# Molecular dynamics unveils multiple-site binding of inhibitors with reduced activity on the surface of dihydrofolate reductase

**DOI:** 10.1101/2024.03.27.586990

**Authors:** Mitsugu Araki, Toru Ekimoto, Kazuhiro Takemura, Shigeyuki Matsumoto, Yunoshin Tamura, Hironori Kokubo, Gert-Jan Bekker, Tsutomu Yamane, Yuta Isaka, Yukari Sagae, Narutoshi Kamiya, Mitsunori Ikeguchi, Yasushi Okuno

## Abstract

The sensitivity to protein inhibitors is altered by modifications or protein mutations, as represented by drug resistance. The mode of stable drug binding to the protein pocket has been experimentally clarified. However, the nature of the binding of inhibitors with reduced sensitivity remains unclear at the atomic level. In this study, we analyzed the thermodynamics and kinetics of inhibitor binding to the surface of wild-type and mutant dihydrofolate reductase (DHFR) using molecular dynamics simulations combined with Markov state modeling. A strong inhibitor of methotrexate (MTX) showed a preference for the active site of wild-type DHFR with minimal binding to unrelated (secondary) sites. Deletion of a side-chain fragment in MTX largely destabilized the active site-bound state, with clear evidence of binding to secondary sites. Similarly, the F31V mutation in DHFR diminished the specificity of MTX binding to the active site. These results reveal the presence of multiple-bound states whose stabilities are comparable to or higher than those of the unbound state, suggesting that a reduction in the binding affinity for the active site significantly elevates the fractions of these states. This study sheds light on the specific drug recognition by proteins and the selectivity of drug binding sites on protein surfaces. (199 words)

## Introduction

Sensitivity to protein inhibitors is reportedly altered by their structural modifications ^1^ or protein mutations ^2^. Decreases in inhibitory activity, particularly drug resistance, is a serious issue in medicine. The molecular mechanisms of the decreased activity have been mostly explained by decreases in binding affinity due to the partial loss of the protein-inhibitor interaction ^3 4 5^. This interpretation is based on a two-state model consisting of a single bound state and an unbound state. However, there is an ensemble of stable and metastable bound states in the solution ^6 7^. The binding modes of drugs stably bound to proteins have been observed using X-ray crystallography, nuclear magnetic resonance spectroscopy, and cryo-electron microscopy. However, since weak protein-drug interactions are still difficult to experimentally capture at the atomic level, the nature of the binding of drugs with decreased sensitivity to their target proteins is not fully understood.

Molecular dynamics (MD) simulations are capable of detecting weak protein-drug interactions at the atomic level. Conventional MD simulations starting from the dissociation state capture only some of the total bound states because of limitations in simulation time (i.e., several microseconds on standard high-performance computers). In contrast, MD simulations with enhanced sampling techniques, such as GaMD ^8^, gREST/REUS ^9^, and McMD ^10^, can exhaustively and efficiently explore metastable and stable drug binding conformations, characterizing their thermodynamic stabilities. Furthermore, recent developments in Markov state modeling (MSM) that integrate multiple conventional MD data have enabled the estimation of the kinetic network among the unbound, stable, and metastable bound states ^11 12^. This has enabled a precise description of drug binding on the protein surface ^13 14^. However, because the huge computational cost (>100 μs) is generally required per protein-drug (inhibitor) pair to build a statistically-reliable model, only a few studies have addressed the effects of drug modifications or protein mutations on both stable and metastable bound states ^15 16^.

In this study, we analyzed the effects of drug modifications or protein mutations on the thermodynamics and kinetics of the binding of an inhibitor to the surface of dihydrofolate reductase (DHFR) using MD simulations combined with MSM. First, exhaustive MD simulations starting from the unbound and inhibitor-bound states were performed for three protein-inhibitor pairs (Fig. 1): (i) wild-type DHFR (DHFR^WT^) and its substrate analogue methotrexate (MTX), (ii) DHFR^WT^ and a MTX analogue lacking the (p-aminobenzoyl)glutamate side-chain (DAM; 2,4-diamino-6,7-dimethylpteridine), and (iii) F31V mutant DHFR (DHFR^F31V^) and MTX. Next, the simulation data acquired for each pair were analyzed using an MSM workflow that algorithmically identified stable and metastable inhibitor binding states on the protein surface. Association/dissociation rate constants and binding stability were estimated for each state. Inhibitor binding to multiple sites, in addition to the active site, was observed during MD simulations for all pairs. Our simulations suggested that, while the strong binder MTX preferred the active site of DHFR^WT^, binding to other sites (secondary sites) was evident by the F31V mutation on DHFR or deletion of the side-chain of MTX. Furthermore, elevated fractions of secondary-site bound states significantly decrease the population of active site-bound states closely associated with inhibitory activity. The details of the binding behaviors of the strong and weak binders obtained in this study provide deep insight into the specific drug recognition of proteins, and the binding site selectivity of drugs on protein surfaces.

**Fig. 1.**
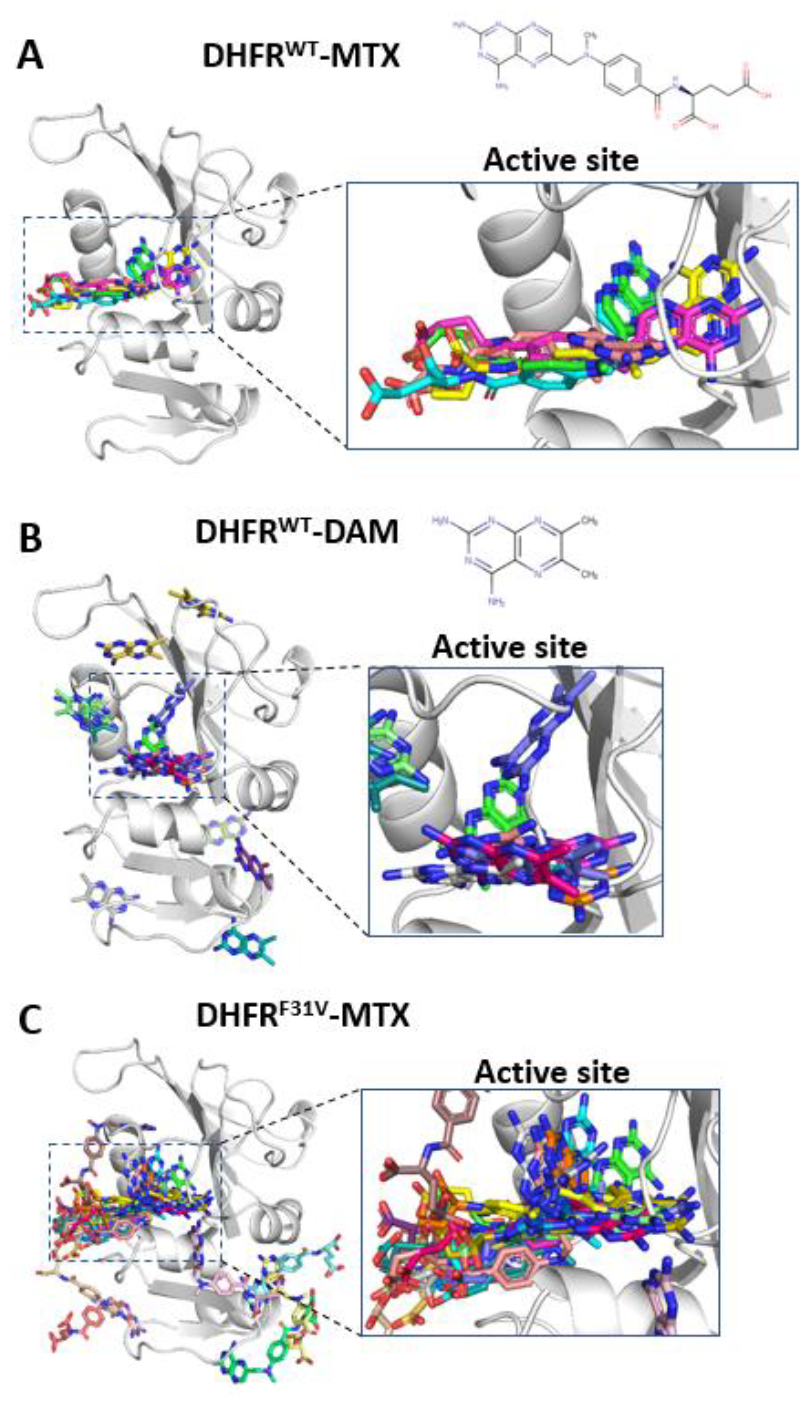
Principal inhibitor binding sites on the DHFR surface. Binding poses of (A) MTX on DHFR^WT^, (B) DAM on DHFR^WT^, and (C) MTX on DHFR^F31V^. The chemical structures of MTX and DAM are shown at the top of A and B, respectively. The protein backbone is depicted with gray ribbons, and representative 5 (A), 18 (B), and 25 (C) inhibitor binding poses are shown with colored sticks, which correspond to the cluster centers of metastable states. The right panels in A-C show an enlarged view of the active site of DHFR.

## Experimental section

### Model systems and force fields

We modeled the complex structure of DHFR and the DAM and MTX inhibitors. based on the cocrystal structure of *Escherichia coli* DHFR in complex with MTX (PDB ID: 1RG7). Missing side chains were modeled and refined using the structure preparation module in the Molecular Operation Environment (MOE) program ^17^, and the dominant protonation state at pH 7.0 was assigned to titratable residues. The F31V mutation was introduced into the structure of wild-type DHFR using the structure preparation module of MOE. To model the initial structure of the unbound state, a DAM or MTX molecule was randomly placed around the protein and away from the binding site (> 15 Å) by translating the bound inhibitor into a cocrystal structure.

The inhibitors were protonated to give net charges of 0 (DAM) or -2 (MTX), reflecting the dominant protonation states at neutral pH. GAMESS was used to optimize the structure of each inhibitor and calculate its electrostatic potential at the HF/6-31G* level ^18^, after which the atomic partial charges were obtained via the restrained electrostatic potential (RESP) approach ^19^. Other potential parameters of the inhibitors were obtained by the general AMBER force field (GAFF) ^20^ using antechamber module of AMBER Tools 12. The AMBER ff99SBILDN force field ^21^ was used for proteins and ions, whereas water was modeled with the TIP3P potential ^22^. Approximately 9,800 water molecules were placed around the protein model in an 69×69×69 Å^3^ cubic box. In addition, approximately 50 sodium and chloride ions (corresponding to 100 mM NaCl) were added to the simulation box to neutralize all systems.

### MD simulations

MD simulations with periodic boundary conditions were performed using the GROMACS 4.6.5 program ^23^. Electrostatic interactions were handled using the particle mesh Ewald method ^24^ with a cutoff radius of 11 Å, and van der Waals interactions were gradually cut off at 10 Å. The P-LINCS algorithm ^25^ was employed to constrain all the bond lengths at their equilibrium values. After energy minimization, each system was equilibrated for 100 ps in a constant number of molecules, volume, and temperature (NVT) ensemble, followed by an MD run of 100 ps in a constant number of molecules, pressure, and temperature (NPT) ensemble with positional restraints applied to protein heavy atoms. The production runs were conducted under NPT conditions without positional restraints. The temperature was maintained at 298 K using a Nose-Hoover thermostat ^26 27^, and a Berendsen barostat ^28^ was used to maintain the pressure at 1 bar. The temperature and pressure time constants were set to 0.3 and 1 ps, respectively. A time step of 2 fs was used in all MD runs. In total, 326, 346, and 347 independent production runs of 100 ns (with different initial atomic velocities) starting from the unbound state were performed for the DHFR^WT^-DAM, DHFR^WT^-MTX, and DHFR^F31V^-MTX systems, respectively. To adequately sample the binding events for the active site, we used an iterative sampling protocol, with three rounds of MD simulations (Fig. S1). In the second round, 260, 400, and 440 different binding structures were extracted for the DHFR^WT^-DAM, DHFR^WT^-MTX, and DHFR^F31V^-MTX systems, respectively, using k-means clustering of the first round of MD trajectories. Simulations of 100 ns were restarted from snapshots corresponding to these states. In the third round, 474 and 672 100-ns simulations for the DHFR^WT^-MTX and DHFR^F31V^-MTX systems, respectively, were restarted from different inhibitor-bound states extracted from the second round of MD trajectories in the same manner. We also performed 50, 200, and 200 independent production runs of 100 ns (with different atomic velocities) starting from the inhibitor-bound state observed in the cocrystal structure (PDB ID: 1RG7) for the DHFR^WT^-DAM, DHFR^WT^-MTX, and DHFR^F31V^-MTX systems, respectively. Subsequently, we extracted the inhibitor-bound states with the 200, 300, and 300 highest root-mean-square deviations (RMSD) from the crystallographic pose of the pteridine moiety from the MD trajectories of the DHFR^WT^-DAM, DHFR^WT^-MTX, and DHFR^F31V^-MTX systems. Additional 100 ns simulations were restarted from snapshots corresponding to these states. The total simulation time was 83.6, 172.0, and 195.9 μs for the DHFR^WT^-DAM, DHFR^WT^-MTX, and DHFR^F31V^-MTX systems, respectively.

### MSM analysis

The aggregated trajectories were analyzed using an MSM-based protocol, as described in Fig. S2 to quantitatively describe the DHFR-inhibitor binding and unbinding processes. PyEMMA ver.2.X ^12^ (http://pyemma.org) was used to construct the MSM from all obtained trajectories. The nearest-neighbor heavy-atom contacts between DHFR residues (159 residues) and the inhibitor were calculated with a cutoff of 5 Å, resulting in a 159-dimentional binary feature vector per trajectory frame. Time-lagged independent component analysis (TICA) ^29-30^ with a lag time of 10 ns was used for dimension reduction, which projected the input feature vectors onto the 53, 80, and 84 slowest TICA components for the DHFR^WT^-DAM, DHFR^WT^-MTX, and DHFR^F31V^-MTX systems, respectively, representing 95% cumulative kinetic variance. The snapshots on the TICA projection map were then clustered into 765, 1300, and 1400 clusters for the DHFR^WT^-DAM, DHFR^WT^-MTX, and DHFR^F31V^-MTX systems, respectively, using k-means clustering, where the number of clusters, *N*_*c*_, was set to √*N*_*f*_, and *N*_*f*_ is the total number of snapshots in the input trajectories. To obtain the appropriate lag time, microstate MSMs were constructed at different lag times (Fig. S3-S5). For all systems, the implied time-scales converged beyond 30 ns, which was consistently used for further analyses. To enable the sensitive detection of differences in the protein-inhibitor binding process dependent on inhibitor modification or protein mutation, macrostates corresponding to stable/metastable states of the DHFR-inhibitor complex were determined as follows. First, the stationary population of all microstates (the number of microstates equals *N*_*c*_) was computed, and microstates with populations higher than 1 / *N*_*c*_ were extracted (*i*.*e*. 174-188 of 765 microstates for the DHFR^WT^-DAM system, 250-273 of 1300 microstates for the DHFR^WT^-MTX system, and 372-380 of 1400 microstates for the DHFR^F31V^-MTX system). Next, clustering of these high-population microstates into an optimal number of macrostates was performed on TICA projection data using the X-means clustering method ^31^, resulting in 8-23, 3-11, and 26-39 meta-stable states with populations of > 0.5% for the DHFR^WT^-DAM, DHFR^WT^-MTX, and DHFR^F31V^-MTX systems, respectively. The association rate constant (*k*_*on*_) and the dissociation rate constant (*k*_*off*_) for each metastable state were calculated as

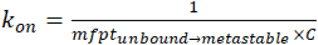

and

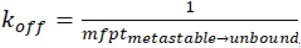, respectively, where *mfpt*_*unbound→metastable*_ is the mean first passage time (MFPT) from the unbound to a metastable state, *mfpt*_*metastable→unbound*_ is MFPT from a metastable to the unbound state, and *C* is the inhibitor (protein) concentration ^12^. Macroscopic k_on_, k_off_, and binding free energy (ΔG) values were estimated according to three models with different definitions of the bound state or the unbound state (Fig. 2 and Table S1). After the metastable states assigned to the bound or unbound state were merged, k_on_ and k_off_ were estimated according to the aforementioned equations based on MFPT, and *ΔG* was calculated according to, where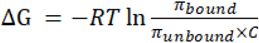, where *R* is the gas constant, *T* is the absolute temperature, *C* is the inhibitor (protein) concentration, and π_bound_ and π_unbound_ are the stationary populations of the bound and unbound state, respectively^12^. To estimate the computational errors of *ΔG, k*_*on*_, and *k*_*off*_, three independent MSM analyses (from k-means clustering to *ΔG, k*_*on*_, and *k*_*off*_ calculations) were performed for each system.

**Fig. 2.**
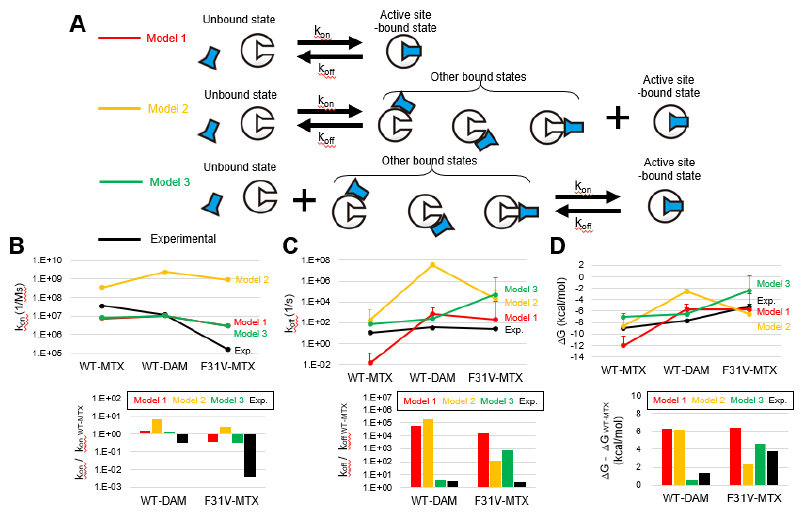
Thermodynamic and kinetic parameters of the DHFR-inhibitor binding process. (A) three models of DHFR-inhibitor binding: (1) from the unbound state to the active site-bound state, (2) from the unbound state to all the bound states, and (3) from the unbound state and all the other bound states to the active site-bound state. (B) The on-rate constant (k_on_), (C) off-rate constant (k_off_), and (D) binding free energy (ΔG) of the DHFR^WT^-MTX, DHFR^WT^-DAM, and DHFR^F31V^-MTX pairs. These values were estimated for each of the three models. The k_on_ and ΔG values were calculated from MSM analyses of binding and unbinding simulation data. After ΔG was corrected by a free energy difference upon protonation of Asp27, k_off_ was calculated using k_on_ and ΔG values (See “Experimental Section” for details). Relative values against DHFR^WT^-MTX are shown in the bottom.

### Estimation of the effect of Asp27 protonation

The binding kinetics of the DHFR^WT^-DAM, DHFR^WT^-MTX, and DHFR^F31V^-MTX pairs were experimentally measured as described by Benkovic et al. ^32^. This measurement was performed at pH 6.0, at which Asp27 of DHFR in the unbound state exists predominantly in the protonated state (pKa=6.5) ^33^. Based on the macroscopic *k*_*on*_, *k*_*off*_, and *ΔG* values estimated from binding and unbinding simulations using DHFR with Asp27 in the deprotonated state, those for DHFR with Asp27 in the protonated state were calculated as follows (Fig. S6 and Table S1). First, we hypothesized that *k*_*on*_ does not change upon the protonation of Asp27 because this residue is located deep within the active site, considering that its protonation/deprotonation would not affect the binding process of the inhibitors. Next, *ΔG* was corrected using a change in the binding free energy upon Asp27 protonation, which was estimated by free energy perturbation (FEP) calculations. For this purpose, we applied MutationFEP, an alchemical FEP method that calculates mutation-induced changes in protein-ligand binding free energy ^34^. Each of the unbound and metastable states extracted from the MSM analysis was set to the initial structure and free energy simulations (27λ × 5 ns) were performed as described previously ^34^. The resulting free energy changes for all states are summarized in Tables S2-S4. Using these values, the changes in *ΔG* were estimated for each model, as shown in Table S1. Finally, using these *k*_*on*_ and modified *ΔG* values, *k*_*off*_ was calculated according to *ΔG* = -*RT*ln (*k*_*on*_ / *k*_*off*_).

## Result and Discussion

### Kinetics and thermodynamics of DHFR-inhibitor binding estimated from MD simulations combined with MSM

Fluorescence spectroscopy has shown that DHFR inhibitors, such as DAM and MTX, bind to the active site of DHFR through a two-step process ^32^. First, these ligands enter the active site (phase 1), and an intermolecular salt bridge is formed via proton transfer between Asp27 of DHFR and the pteridine moiety of the inhibitors within pocket ^35^ (phase 2). The experimental measurement of the binding kinetics for phase 1 showed that the association rate and binding affinity of DAM and MTX with DHFR^WT^ were significantly higher than those of MTX with DHFR^F31V 32^, in which a conserved residue at the active site, Phe31, was replaced with valine. In this study, we focused on phase 1 of the DHFR-inhibitor binding process and performed binding and unbinding simulations for each of the DHFR^WT^ - MTX, DHFR^WT^ - DAM, and DHFR^F31V^ - MTX pairs.

When approximately 300 independent 100 ns MD simulations were performed from the unbound state, 31, 33, and 22 inhibitor binding events to the active site were captured for the DHFR^WT^ - MTX, DHFR^WT^ - DAM, and DHFR^F31V^ - MTX systems, respectively. After performing additional simulations to enrich the conformational sampling of the bound inhibitor, all MD trajectories acquired for each pair were analyzed using an MSM analysis workflow that algorithmically assigned metastable states of the protein-ligand complex (Fig. S2). As a result, multiple inhibitor-bound states were identified for all the pairs, including the crystallographic binding mode ^36^ (Fig. 1).

The number of the MSM-defined metastable states of the DHFR-inhibitor complex was 6 ± 3 for DHFR^WT^ - MTX, 16 ± 7 for DHFR^WT^ - DAM, and 26 ± 6 for DHFR^F31V^ - MTX. The on-rate constant (k_on_), off-rate constant (k_off_), and binding free energy (ΔG) were estimated according to three models with different definitions of the bound state or the unbound state (Fig. 2A): (1) binding to the active site from the unbound state, (2) binding to all the sites on the DHFR surface from the unbound state, and (3) binding to the active site from the other states. While model (1), which is most generally used for analyzing protein-ligand binding, showed smaller deviations from the experimentally measured k_on_ and k_off_ values (Fig. 2B and 2C) ^32^, this model provided comparable ΔG values for the DHFR^WT^ - DAM and DHFR^F31V^ – MTX pairs (Fig. 2D). In contrast, model (3) best reproduced the relative relationship between the experimental values at the point where MTX binding to DHFR^F31V^ exhibited the lowest on-rate and binding affinity among the three pairs (Fig. 2B and 2D). The results may suggest that, while the inhibitors bound tightly to the active site efficiently quenched the fluorescence of tryptophans in DHFR ^32^, the other bound states did not have sufficient quenching ability, and these states were experimentally indistinguishable from the unbound state.

### Microscopic pictures of inhibitor binding processes

Microscopic images of the inhibitor binding process on the DHFR surface were estimated by calculating the association/dissociation rates and stability of each metastable state of the DHFR-inhibitor complex (Fig. 3-5). The A smaller molecular weight inhibitor, DAM, accessed the active site and multiple secondary sites with comparable association rate constants of ∼10^7^ (M^-1^s^-1^) and binding stabilities (Fig. 3). One of the estimated metastable states (metastable state index = 1), which coincides with the binding mode of the pteridine moiety observed in the DHFR^WT^ – MTX cocrystal structure ^36^, was isolated from other metastable states in the free energy landscape of inhibitor binding (Fig. 3A). The finding suggests the existence of an energy barrier to the transition to this state. This state provided the smallest association/dissociation rates [i.e. k_on_ of 1.1± 0.0 ×10^7^ (M^-1^s^-1^) and k_off_ of 7.2± 0.3 ×10^5^ (s^-1^) obtained from three independent MSM analyses] among all the metastable states (Fig. 3B and 3C) although its stability was not significantly higher than those of the other states (Fig. 3D). In a main binding pathway the ligand binds to the active site via “encounter complexes” or “doorway states” ^37^ (e.g. metastable state index = 6 and 11), in which the ligand weakly bound to the entrance of the active site (Fig. 3A).

**Fig. 3.**
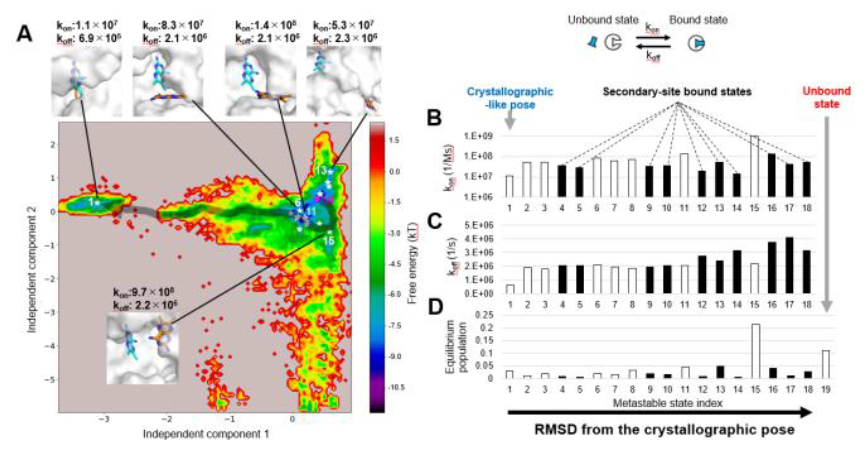
Thermodynamics and kinetics of DHFR^WT^-DAM binding. (A) The free energy landscape projected onto the first and second time-lagged independent components calculated from TICA. Eighteen metastable states of the DHFR-inhibitor complex with populations > 0.5% and the dissociation state are indicated by white stars and magenta diamonds, respectively. (B-D) The association rate constant from the unbound state (k_on_) (B), dissociation rate constant to the unbound state (k_off_) (C), and equilibrium population (D) are plotted for each metastable state. A state with lower and higher root-mean-square deviation (RMSD) from the crystallographic pose is labeled with lower and higher state indices, respectively. Intermediate states in binding to the active site (open bars) were extracted from a flux analysis (Table S5). The other states are defined as secondary-site bound states (filled bars). State 19 corresponds to the unbound state. In (A), conformations of representative metastable states (orange sticks) on DHFR (white surfaces) are shown alongside the crystallographic pose (cyan sticks), together with their k_on_ and k_off_ values. A representative binding pathway to the active site via high flux states is indicated by a gray arrow.

MTX is a larger molecular weight inhibitor in which a bulky acidic side-chain fragment is added to the DAM moiety. This inhibitor did not bind to most of the secondary sites detected in the DAM simulations (Fig. 4). Metastable states, excluding the metastable state index = 4, could be roughly categorized into only two types (Fig. 4A): the doorway states defined above (metastable state index = 3 and 5) and the active site-bound state corresponding to the crystallographic binding mode ^36^ (metastable state index = 1, 2), suggesting that a single minimum free energy path is formed in the DHFR binding of MTX, whose chemical structure is very similar to that of its folate substrate. As well as the simulation result of DAM, the association/dissociation rates for this state [i.e. k_on_ of 7.6 ± 0.5 ×10^6^ (M^-1^s^-1^) and k_off_ of 2.9 ± 0.1 ×10^4^ (s^-1^)] are smaller than those for any other bound states (Fig. 4B and 4C). These results indicate that the addition of the side-chain fragment to the DAM moiety increased the binding specificity for the active site by reducing the association rate and binding stability for secondary sites with small pocket sizes.

**Fig. 4.**
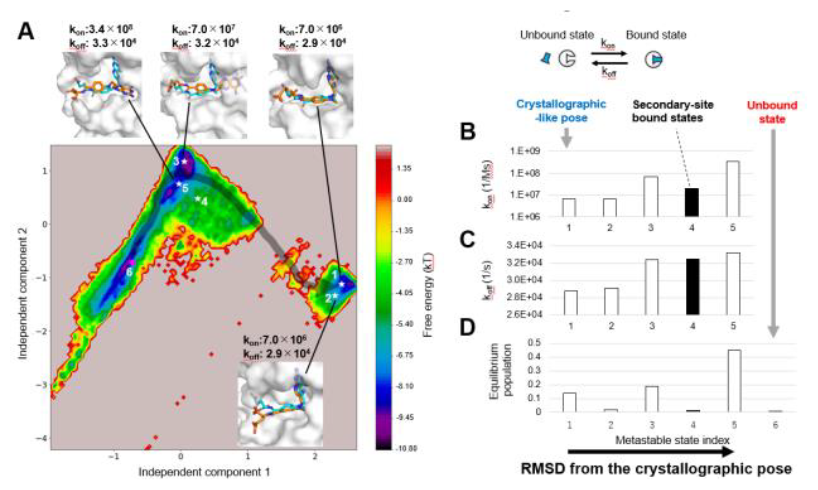
Thermodynamics and kinetics of DHFR^WT^-MTX binding. (A) The free energy landscape projected onto the first and second time-lagged independent components calculated from TICA. Five metastable states of the DHFR-inhibitor complex with populations > 0.5% and the dissociation state are indicated by white stars and magenta diamonds, respectively. (B-D) The association rate constant from the unbound state (k_on_) (B), dissociation rate constant to the unbound state (k_off_) (C), and equilibrium population (D) are plotted for each metastable state, where a state with lower and higher root-mean-square deviation (RMSD) from the crystallographic pose are labeled with lower and higher state indices, respectively. Intermediate states in binding to the active site (open bars) were extracted from a flux analysis (Table S6). The other state is defined as secondary-site bound states (filled bars). State 6 corresponds to the unbound state. In (A), conformations of representative metastable states (orange sticks) on DHFR (white surfaces) are shown alongside the crystallographic pose (cyan sticks), together with their k_on_ and k_off_ values. A representative binding pathway to the active site via high flux states is indicated by a gray arrow.

The number of metastable MTX binding states was remarkably increased in the F31V mutant compared to the wild-type (Fig. 5). This is attributed to the fact that k_on_ for the active site was reduced from 7.6 ± 0.5 × 10^6^ to 2.8 ± 0.3 × 10^6^ (M^-1^s^-1^), while those for secondary sites was increased to 10^7^ – 10^8^ (M^-1^s^-1^) by the mutation (Fig. 4B and 5B). Because Phe31 is one of the active site-forming residues ^36^ and its replacement with valine narrowed the shape of the pocket (Fig. S7), the decrease in k_on_ for the active site appears to result from the difficulty in accessing the pocket of the mutant. These results imply that the F31V mutation decreased the binding specificity for the active site by destabilizing the binding pathway to the pocket.

**Fig. 5.**
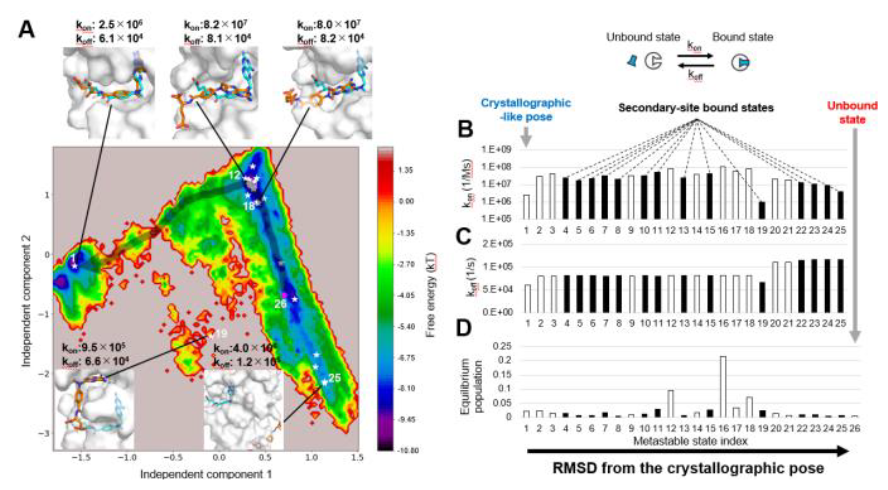
Thermodynamics and kinetics of DHFR^F31V^-MTX binding. (A) The free energy landscape projected onto the first and second time-lagged independent components calculated from TICA. Twenty five metastable states of the DHFR-inhibitor complex with populations of > 0.5% and the dissociation state are indicated by white stars and magenta diamonds, respectively. (B-D) The association rate constant from the unbound state (k_on_) (B), dissociation rate constant to the unbound state (k_off_) (C), and equilibrium population (D) are plotted for each metastable state, where a state with lower and higher root-mean-square deviation (RMSD) from the crystallographic pose are labeled with lower and higher state indices, respectively. Intermediate states in binding to the active site (open bars) were extracted from a flux analysis (Table S7). The other states are defined as secondary-site bound states (filled bars). State 26 corresponds to the unbound state. In (A), conformations of representative metastable states (orange sticks) on DHFR (white surfaces) are shown alongside the crystallographic pose (cyan sticks), together with their k_on_ and k_off_ values. A representative binding pathway to the active site via high flux states is indicated by a gray arrow.

### Effect of DHFR mutation or modification of the inhibitors

Binding and unbinding simulations for all three pairs captured the active site-bound and intermediate states along pathways to this site, but also multiple secondary-site bound states. Many of their stabilities were equivalent to or higher than those of the unbound state (Fig. 3-5). The stability of the active site-bound state was remarkably higher in the DHFR^WT^-MTX pair than in the DHFR^WT^-DAM and DHFR^F31V^-MTX pairs (Table S1). This is consistent with the results of the commonly used binding FEP method, MP-CAFEE ^38^, which directly computesΔG corresponding to model (1) in Fig. 2 (Table S8). The higher binding affinity of MTX than that of DAM for DHFR^WT^ was attributed to additional electrostatic and van der Waals interactions with the side-chain fragment introduced to the DAM moiety (Fig. 6A and Table S8). The F31V mutation, which replaces a bulky side-chain with a smaller one, reduced the electrostatic interactions with MTX by slightly shifting the bound MTX toward the mutated residue (Fig. 6B and Table S8).

**Fig. 6.**
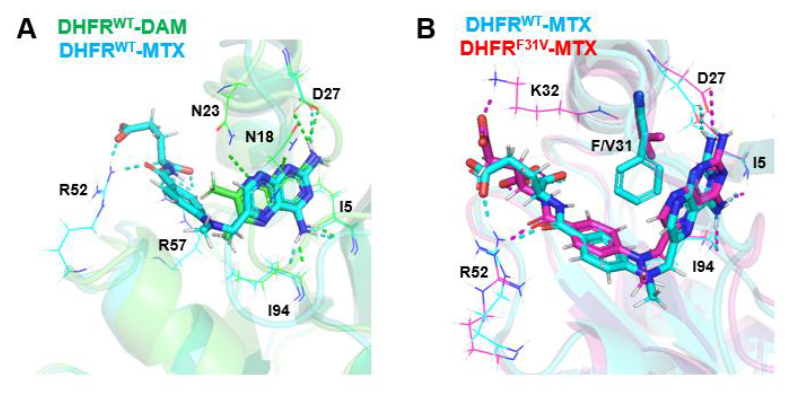
Structural comparison of the DHFR^WT^-DAM, DHFR^WT^-MTX, and DHFR^F31V^-MTX complexes. A complex structure closest to the cluster center of metastable state index = 1 was extracted and compared between the DHFR^WT^-DAM and DHFR^WT^-MTX complexes (left) and between the DHFR^WT^-MTX and DHFR^F31V^-MTX complexes (right). The protein backbone is represented by a transparent ribbon diagram. The inhibitors and side chains of F/V31 are depicted as thick sticks, while those of the other residues are depicted as thin sticks (C, green/cyan/magenta; N, blue; O, red). DHFR-inhibitor hydrogen bonds are represented by dotted lines.

The population of active site-bound states can be regarded as a physicochemical parameter closely associated with the activity of protein inhibitors. In strong binders, such as DHFR^WT^-MTX, because the fractions of secondary-site bound states are negligibly small, the population of the active site-bound state could be correctly estimated by FEP methods, which assume only two states of the unbound and bound states. In contrast, in weak binders, such as DHFR^WT^-DAM and DHFR^F31V^-MTX, FEP may overestimate the population of the active site-bound state because of the significantly increased fractions of the secondary-site bound states (Fig. 7). Therefore, accurate estimation of the inhibitory activity of a weak binder would require evaluation of the stability of the active site-bound state relative to the dissociated state and secondary-site bound states. As demonstrated in this study, a combination of binding simulations starting from the unbound state and MSM is useful for the assessment of protein-inhibitor associations, including secondary binding sites. Alternatively, generalized ensemble MD simulations, such as accelerated-MD ^8^, gREST ^9^, and McMD ^10^, may be effective for efficiently exploring inhibitor binding sites on the protein surface and estimating the binding stability for each site.

**Fig. 7.**
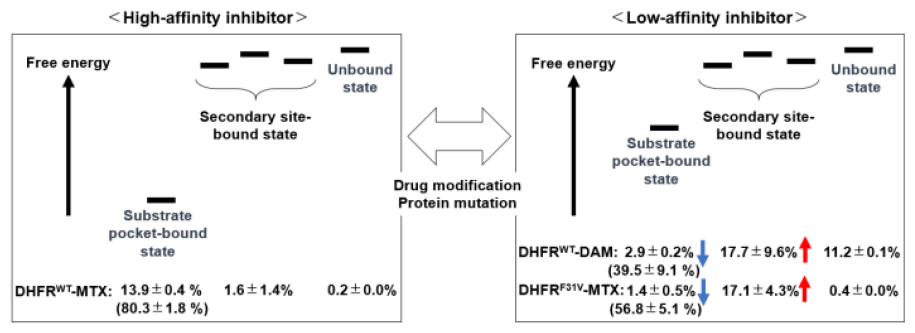
Equilibrium distribution of high/low-affinity inhibitors. The populations of the substrate pocket-bound state, secondary-site bound states (corresponding to filled bars in Fig. 3-5), and unbound state are indicated. Their remarkable increases and decreases are highlighted with red and blue arrows, respectively. Sum of the populations of the substrate pocket-bound state and intermediate states along pathways to the pocket (corresponding to open bars in Fig. 3-5) is indicated in parentheses. The values are presented as the mean ± SD of three independent MSM analyses.

## Conclusion

In this study, we analyzed the effects of protein mutations and drug modifications on inhibitor binding to the DHFR surface using MD simulations combined with MSM. The weak inhibitor, DAM, exhibited low binding affinity for the substrate pocket, identifying binding to multiple secondary sites. The addition of a side-chain fragment to DAM, which increased its inhibitory activity by more than 1000-fold, resulted in a much higher binding affinity for the substrate pocket and significantly decreased the fractions of secondary-site bound states. However, the F31V mutation that occurred in the substrate pocket reduced the binding affinity of MTX to this pocket, with a considerable elevation in the fractions of secondary-site bound states. Traditional binding free energy computation methods (*e*.*g*. FEP methods), which assume a two-state model consisting of a single bound state and an unbound state, have been employed to assess the activity of protein inhibitors. However, since the aforementioned results demonstrated that inhibitors with reduced binding affinity for the active site prefer to bind to other sites on the protein surface, binding of these inhibitors might not be fully described by these traditional methods, highlighting the importance of considering binding to these secondary sites.

## ASSOCIATED CONTENT

### Supporting Information

Supporting Information includes the following data:

1. Iterative sampling protocol for DHFR-inhibitor binding/unbinding simulations.
2. Flowchart of the MSM analysis protocol.
3. Lag time dependence of the 10 slowest implied time scales for the DHFR^WT^-MTX, DHFR^WT^-DAM, and DHFR^F31V^-MTX systems.
4. A proposed DHFR-inhibitor binding scheme.
5. Structural comparison of the DHFR^WT^-MTX and DHFR^F31V^-MTX complexes.
6. Kinetic and thermodynamic parameters of inhibitor binding to DHFR with Asp27 in the deprotonated and protonated states.
7. Free energy changes of the DHFR^WT^-MTX, DHFR^WT^-DAM, and DHFR^F31V^-MTX complexes upon protonation of Asp27.
8. Contributions of each metastable state of the DHFR^WT^-MTX, DHFR^WT^-DAM, and DHFR^F31V^-MTX complexes in a network of fluxes from the unbound state to the active site-bound state.
9. DHFR^WT^-MTX, DHFR^WT^-DAM, and DHFR^F31V^-MTX binding free energies (ΔG) estimated by the MP-CAFEE method.

## AUTHOR INFORMATION

### Author Contributions

The manuscript was written through contributions of all authors. /All authors have given approval to the final version of the manuscript. +These authors contributed equally.

## Notes

The authors declare no competing interests.

## ACKNOWLEDGMENT

This study was supported by the Ministry of Education, Culture, Sports, Science and Technology (MEXT, Japan) as “Program for Promoting Researches on the Supercomputer Fugaku” (Simulation- and AI-driven next-generation medicine and drug discovery based on “Fugaku,” JPMXP1020230120) (to Y.O), the K super-computer-based drug discovery project by Biogrid pharma consortium (to Y.O),and a Japan Society for the Promotion of Science (JSPS) KAKENHI Grant (No. JP21K06510) (to M.A). The simulations were performed on the K computer and Fugaku super-computer provided by the RIKEN Center for Computational Science through the HPCI System Research Project (project IDs: hp170036, hp180011, hp190020, hp200011, and hp230216).

## ABBREVIATIONS

DAM: 2,4-diamino-6,7-dimethylpteridine
DHFR: dihydrofolate reductase
MD: molecular dynamics
MSM: Markov state model
MTX: methotrexate; RMSD, root-mean-square deviation

## REFERENCES

1. Dörwald, F. Z. Lead Optimization for Medicinal Chemists: Pharmacokinetic Properties of Functional Groups and Organic Compounds, 2012.

2. Drug Resistance in Bacteria, Fungi, Malaria, and Cancer, Springer, 2017.

3. Kobayashi, S.; Boggon, T. J.; Dayaram, T.; Jänne, P. A.; Kocher, O.; Meyerson, M.; Johnson, B. E.; Eck, M. J.; Tenen, D. G.; Halmos, B. EGFR Mutation and Resistance of Non-small-Cell Lung Cancer to Gefitinib. N. Engl. J. Med. 2005, 352, 786–792.

4. Okada, K.; Araki, M.; Sakashita, T.; Ma, B.; Kanada, R.; Yanagitani, N.; Horiike, A.; Koike, S.; Oh-hara, T.; Watanabe, K.; Tamai, K.; Maemondo, M.; Nishio, M.; Ishikawa, T.; Okuno, Y.; Fujita, N.; Katayama, R. Prediction of ALK Mutations Mediating ALK-TKIs Resistance and Drug Re-purposing to Overcome the Resistance. EBiomedicine. 2019, 41, 105–119.

5. Solomon, B. J.; Tan, L.; Lin, J. J.; Wong, S. Q.; Hollizeck, S.; Ebata, K.; Tuch, B. B.; Yoda, S.; Gainor, J. F.; Sequist, L. V.; Oxnard, G. R.; Gautschi, O.; Drilon, A.; Subbiah, V.; Khoo, C.; Zhu, E. Y.; Nguyen, M.; Henry, D.; Condroski, K. R.; Kolakowski, G. R.; Gomez, E.; Ballard, J.; Metcalf, A. T.; Blake, J. F.; Dawson, S. J.; Blosser, W.; Stancato, L. F.; Brandhuber, B. J.; Andrews, S.; Robinson, B. G.; Rothenberg, S. M. RET Solvent Front Mutations Mediate Acquired Resistance to Selective RET Inhibition in RET-Driven Malignancies. J. Thorac. Oncol. 2020, 15, 541–549.

6. Pedersen, M. E.; Haegebaert, R. M. S.; Østergaard, J.; Jensen, H. Size-Based Characterization of Adalimumab and TNF-α Interactions Using Flow Induced Dispersion Analysis: Assessment of Avidity-Stabilized Multiple Bound Species. Sci. Rep. 2021, 11, 4754.

7. Zhao, J.; Blayney, A.; Liu, X.; Gandy, L.; Jin, W.; Yan, L.; Ha, J. H.; Canning, A. J.; Connelly, M.; Yang, C.; Liu, X.; Xiao, Y.; Cosgrove, M. S.; Solmaz, S. R.; Zhang, Y.; Ban, D.; Chen, J.; Loh, S. N.; Wang, C. EGCG Binds Intrinsically Disordered N-Terminal Domain of p53 and Disrupts p53-MDM2 Interaction. Nat. Commun. 2021, 12, 986.

8. Miao, Y.; Bhattarai, A.; Wang, J. Ligand Gaussian Accelerated Molecular Dynamics (LiGaMD): Characterization of Ligand Binding Thermodynamics and Kinetics. J. Chem. Theor. Comput. 2020, 16, 5526–5547.

9. Re, S.; Oshima, H.; Kasahara, K.; Kamiya, M.; Sugita, Y. Encounter Complexes and Hidden Poses of Kinase-Inhibitor Binding on the Free-Energy Landscape. Proc. Natl. Acad. Sci. U. S. A. 2019, 116, 18404–18409.

10. Bekker, G. J.; Araki, M.; Oshima, K.; Okuno, Y.; Kamiya, N. Exhaustive Search of the Configurational Space of Heat-Shock Protein 90 With Its Inhibitor by Multicanonical Molecular Dynamics Based Dynamic Docking. J. Comp. Chem. 2020, 41, 1606–1615.

11. Lawrenz, M.; Shukla, D.; Pande, V. S. Cloud Computing Approaches for Prediction of Ligand Binding Poses and Pathways. Sci. Rep. 2015, 5, 7918.

12. Scherer, M. K.; Trendelkamp-Schroer, B.; Paul, F.; Pérez-Hernández, G.; Hoffmann, M.; Plattner, N.; Wehmeyer, C.; Prinz, J. H.; Noé, F. PyEMMA 2: A Software Package for Estimation, Validation, and Analysis of Markov Models. J. Chem. Theor. Comput. 2015, 11, 5525–5542.

13. Plattner, N.; Noé, F. Protein Conformational Plasticity and Complex Ligand-Binding Kinetics Explored by Atomistic Simulations and Markov Models. Nat. Commun. 2015, 6, 7653.

14. Koneru, J. K.; Sinha, S.; Mondal, J. Molecular Dynamics Simulations Elucidate Oligosaccharide Recognition Pathways by Galectin-3 at Atomic Resolution. J. Biol. Chem. 2021, 297, 101271.

15. Plattner, N.; Doerr, S.; De Fabritiis, G.; Noé, F. Complete Protein-Protein Association Kinetics in Atomic Detail Revealed by Molecular Dynamics Simulations and Markov Modelling. Nat. Chem. 2017, 9, 1005–1011.

16. Pantsar, T.; Kaiser, P. D.; Kudolo, M.; Forster, M.; Rothbauer, U.; Laufer, S. A. Decisive Role of Water and Protein Dynamics in Residence Time of p38α MAP Kinase Inhibitors. Nat. Commun. 2022, 13, 569.

17. Montreal, Q. C.; Canada, H. Molecular Operating Environment (MOE), 2016.08, Chemical Publishing Computing Group Inc.: 1010 Sherbrooke St. West, Suite #910, 2016, p 3A 2R7.

18. Schmidt, M. W.; Baldridge, K. K.; Boatz, J. A.; Elbert, S. T.; Gordon, M. S.; Jensen, J. H.; Koseki, S.; Matsunaga, N.; Nguyen, K. A.; Su, S.; Windus, T. L.; Dupuis, M.; Montgomery, J. A. General Atomic and Molecular Electronic Structure System. J. Comput. Chem. 1993, 14, 1347–1363.

19. Bayly, C. I.; Cieplak, P.; Cornell, W.; Kollman, P. A. A Well-Behaved Electrostatic Potential Based Method Using Charge Restraints for Deriving Atomic Charges: The RESP Model. J. Phys. Chem. 1993, 97, 10269–10280.

20. Wang, J.; Wolf, R. M.; Caldwell, J. W.; Kollman, P. A.; Case, D. A. Development and Testing of a General Amber Force Field. J. Comput. Chem. 2004, 25, 1157–1174.

21. Lindorff-Larsen, K.; Piana, S.; Palmo, K.; Maragakis, P.; Klepeis, J. L.; Dror, R. O.; Shaw, D. E. Improved Side-Chain Torsion Potentials for the Amber ff99SB Protein Force Field. Proteins. 2010, 78, 1950–1958.

22. Jorgensen, W. L.; Chandrasekhar, J.; Madura, J. D.; Impey, R. W.; Klein, M. L. Comparison of Simple Potential Functions for Simulating Liquid Water. J. Chem. Phys. 1983, 79, 926–935.

23. Hess, B.; Kutzner, C.; van der Spoel, D.; Lindahl, E. GROMACS 4: Algorithms for Highly Efficient, Load-Balanced, and Scalable Molecular Simulation. J. Chem. Theor. Comput. 2008, 4, 435–447.

24. Darden, T.; York, D.; Pedersen, L. Particle Mesh Ewald: An N⋅Log(N) Method for Ewald Sums in Large Systems. J. Chem. Phys. 1993, 98, 10089–10092.

25. Hess, B.; Bekker, H.; Berendsen, H. J. C.; Fraaije, J. G. E. M. Lincs: A Linear Constraint Solver for Molecular Simulations. J. Comput. Chem. 1997, 18, 1463–1472.

26. Nosé, S. A Molecular Dynamics Method for Simulations in the Canonical Ensemble. Mol. Phys. 1984, 52, 255–268.

27. Hoover, W. G. Canonical Dynamics: Equilibrium PhaseSpace Distributions. Phys. Rev. A Gen. Phys. 1985, 31, 1695–1697.

28. Berendsen, H. J. C.; Postma, J. P. M.; van Gunsteren, W. F.; DiNola, A.; Haak, J. R. Molecular Dynamics With Coupling to an External Bath. J. Chem. Phys. 1984, 81, 3684–3690.

29. Pérez-Hernández, G.; Paul, F.; Giorgino, T.; De Fabritiis, G.; Noé, F. Identification of Slow Molecular Order Parameters for Markov Model Construction. J. Chem. Phys. 2013, 139, 015102.

30. Schwantes, C. R.; Pande, V. S. Improvements in Markov State Model Construction Reveal Many Non-native Interactions in the Folding of NTL9. J. Chem. Theor. Comput. 2013, 9, 2000–2009.

31. Ishioka, T. Extended K-Means With an Efficient Estimation of the Number of Clusters. In Intelligent Data Engineering and Automated Learning—IDEAL, 2000, pp 17–22.

32. Taira, K.; Benkovic, S. J. Evaluation of the Importance of Hydrophobic Interactions in Drug Binding to Dihydrofolate Reductase. J. Med. Chem. 1988, 31, 129–137.

33. Fierke, C. A.; Johnson, K. A.; Benkovic, S. J. Construction and Evaluation of the Kinetic Scheme Associated With Dihydrofolate Reductase From Escherichia coli. Biochemistry. 1987, 26, 4085–4092.

34. Ono, F.; Chiba, S.; Isaka, Y.; Matsumoto, S.; Ma, B.; Katayama, R.; Araki, M.; Okuno, Y. Improvement in Predicting Drug Sensitivity Changes Associated With Protein Mutations Using a Molecular Dynamics Based Alchemical Mutation Method. Sci. Rep. 2020, 10, 2161.

35. Schlegel, H. B.; Poe, M.; Hoogsteen, K. Models for the Binding of Methotrexate to Escherichia coli Dihydrofolate Reductase: Direct Effect of Carboxylate of Aspartic Acid 27 Upon Ultraviolet Spectrum of Methotrexate. Mol. Pharmacol. 1981, 20, 154–158.

36. Sawaya, M. R.; Kraut, J. Loop and Subdomain Movements in the Mechanism of Escherichia coli Dihydrofolate Reductase: Crystallographic Evidence. Biochemistry. 1997, 36, 586–603.

37. Mondal, J.; Friesner, R. A.; Berne, B. J. Role of Desolvation in Thermodynamics and Kinetics of Ligand Binding to a Kinase. J. Chem. Theor. Comput. 2014, 10, 5696–5705.

38. Fujitani, H.; Tanida, Y.; Matsuura, A. Massively Parallel Computation of Absolute Binding Free Energy With Well-Equilibrated States. Phys. Rev. E Stat. Nonlin. Soft Matter Phys. 2009, 79, 021914.

