## supplemental figures and tables for "Molecular dynamics unveils multiple-site binding of inhibitors with reduced activity on the surface of dihydrofolate reductase"

#### Contents

**1 Supplementary Figures** ... **3**

**2 Supplementary Tables** ... **11**

**3 Supplementary References** ... **19**

### 1. Supplementary Figures

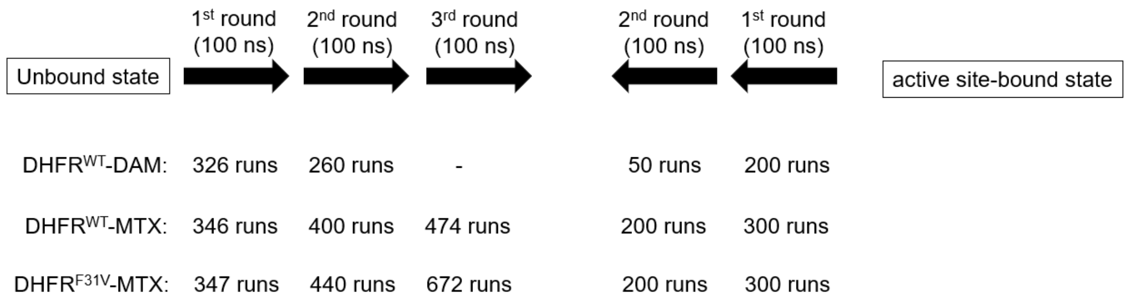


Figure S1 Iterative sampling protocol for dihydrofolate reductase (DHFR)-inhibitor binding/unbinding simulations. The detailed protocol of each round is described in Materials & Methods.

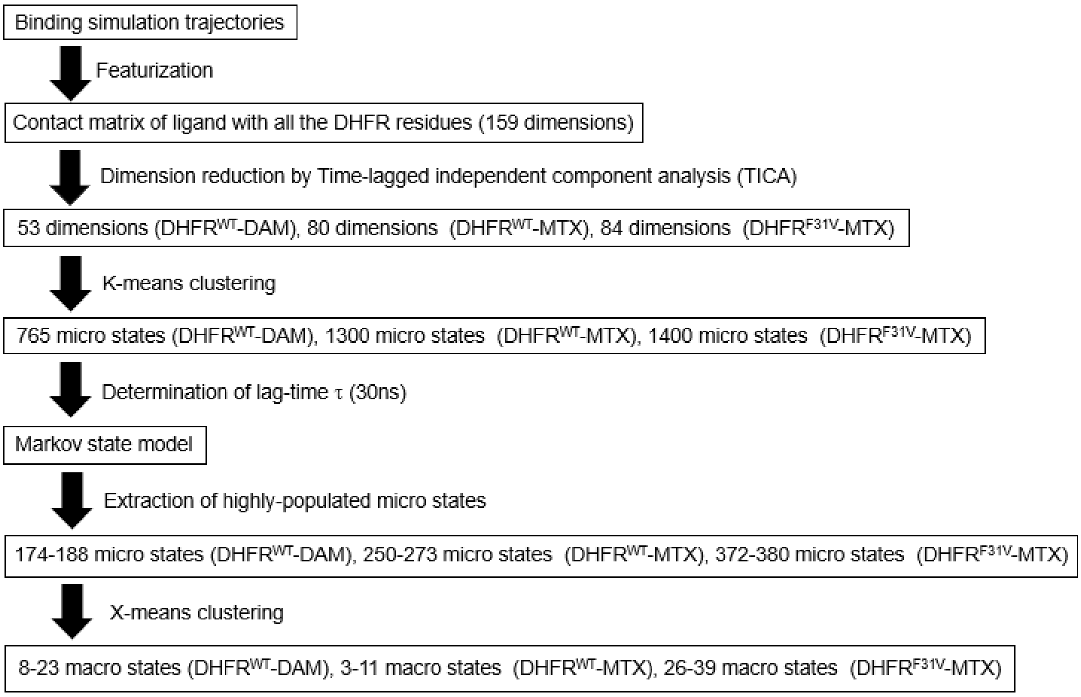


Figure S2. Flowchart of the Markov state modeling (MSM) analysis protocol applied in this study. The detailed protocol of each step is described in Materials & Methods.

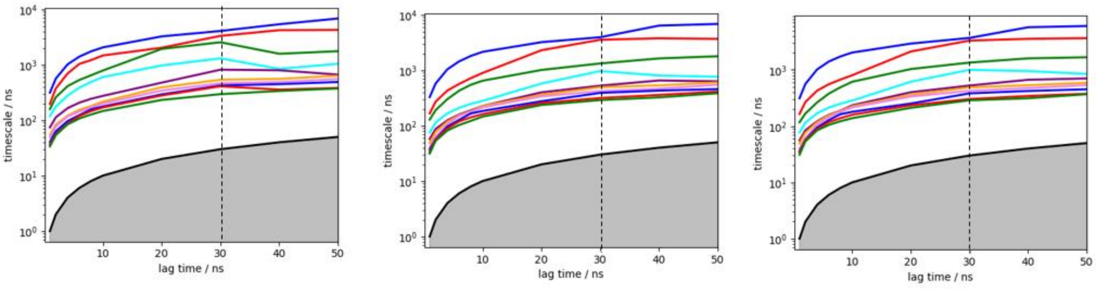


Figure S3 Lag time dependence of the 10 slowest implied time scales for the DHFR$^{WT}$-MTX system. The results of three independent analyses are shown. A lag time of 30 ns was chosen to build the MSM for subsequent analysis, considering that the implied time scales converged at this time.

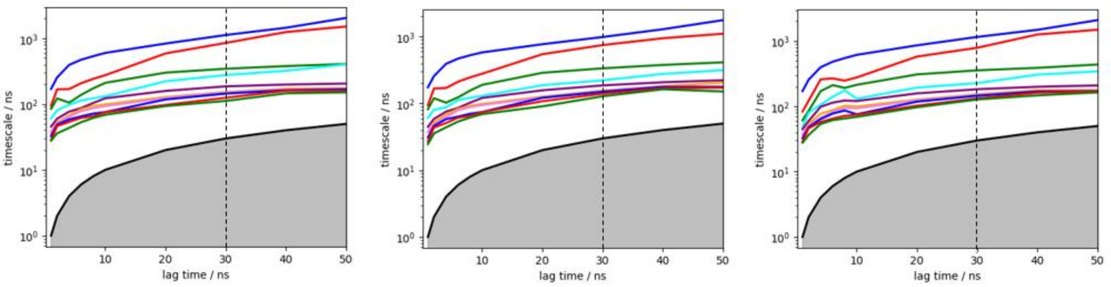


Figure S4 Lag time dependence of the 10 slowest implied time scales for the DHFR$^{WT}$-DAM system. The results of three independent analyses are shown. A lag time of 30 ns was chosen to build the MSM for subsequent analysis, considering that the implied time scales converged at this time.

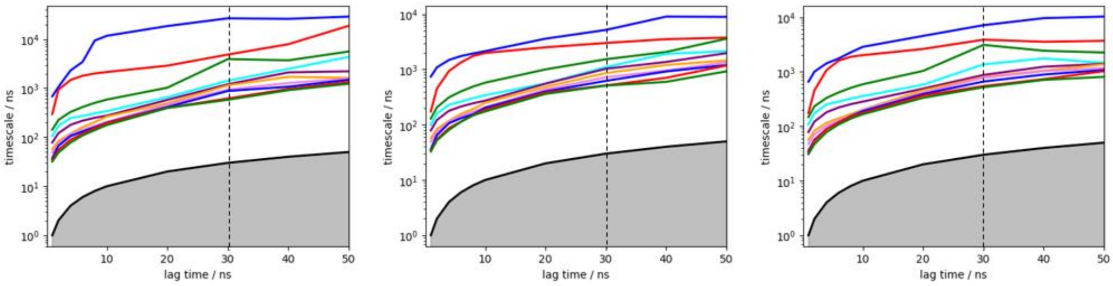


Figure S5 Lag time dependence of the 10 slowest implied time scales for the $DHFR^{F31V}$-MTX system. The results of three independent analyses are shown. A lag time of 30 ns was chosen to build the MSM for subsequent analysis, considering that the implied time scales converged at this time.

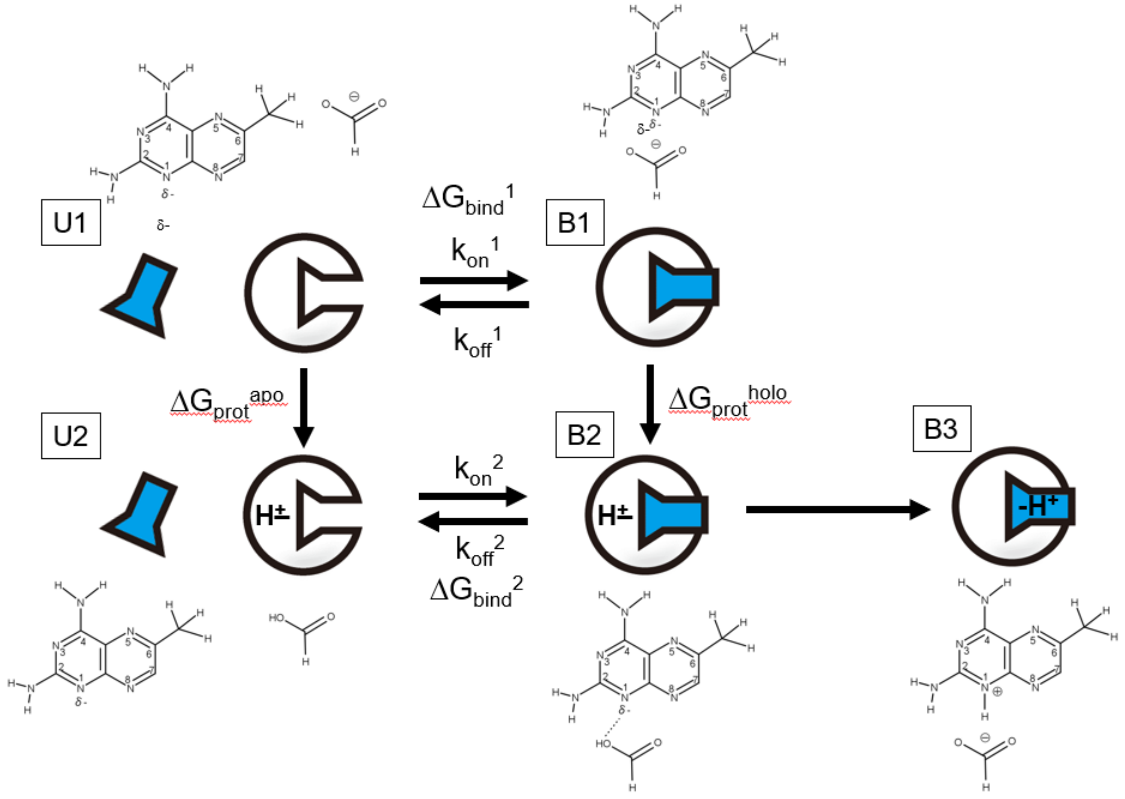


Figure S6 DHFR-MTX (DAM) binding scheme proposed by Benkovic et al [1].

Asp27 of DHFR in the unbound state exists as a mixture of deprotonated (U1) and protonated (U2) states at neutral pH (pKa=6.5). After the inhibitor binds to the active sites of DHFR (B1 and B2), the equilibrium between these two states shifts to a protonated state (B2). Finally, the inhibitor is protonated by proton transfer from Asp27, resulting in the formation of a tight DHFR-inhibitor complex (B3).

In this study, binding and unbinding simulations were performed using DHFR in the deprotonated state (i.e., binding process between U1 and B1) because the dissociation of the inhibitor from DHFR in the protonated state is not expected to occur within ~100 ns in the molecular dynamics simulations because of the formation of an intermolecular hydrogen bond. Thus, the association and dissociation rate constants and binding free energy estimated from these simulations combined with Markov state modeling correspond to. $k_{on}^1$, $k_{off}^1$, and $\Delta G_{bind}^1$, respectively. On the other hand, it is likely that the experimentally measured values (Table 1 of [1]) correspond to $k_{on}^2$, $k_{off}^2$, and $\Delta G_{bind}^2$ because the binding kinetics were measured at pH 6.0, in which Asp27 of DHFR in the unbound state exists dominantly as the protonated state (U2).

To assess the accuracy of our simulations, $k_{on}^2$, $k_{off}^2$, and $\Delta G_{bind}^2$ were estimated using $k_{on}^1$ and $\Delta G_{bind}^1$. First, we hypothesized that $k_{on}^2 = k_{on}^1$ because Asp27 is located deep within the active site, considering that its protonation/deprotonation would not affect the height of the energy barrier for the binding process of the inhibitors. Next, $\Delta G_{bind}^{2}$ was estimated according to $\Delta G_{bind}^{2} = \Delta G_{bind}^{1} + \Delta G_{prot}^{holo} - \Delta G_{prot}^{apo}$, where $\Delta G_{prot}^{holo}$ and $\Delta G_{prot}^{apo}$ are the free energy changes upon protonation of Asp27 in the bound and unbound states, respectively, using MutationFEP [2] . Finally, $k_{off}^{2}$ was estimated according to $k_{off}^{2} = k_{on}^{2} / \exp(-\Delta G_{bind}^{2}/RT)$, where R is the gas constant and T is the absolute temperature.

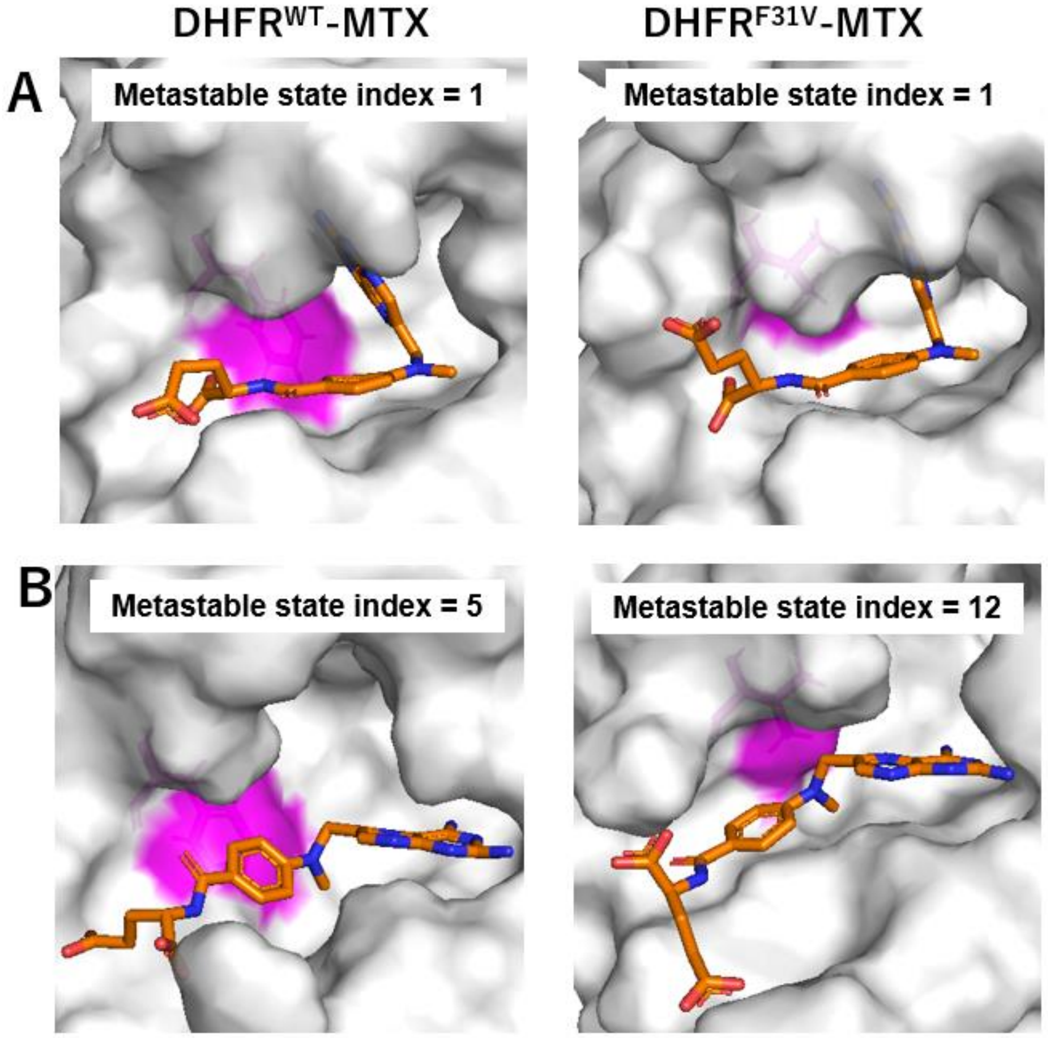


Fig. S7 Structural comparison of the DHFR$^{WT}$-MTX (left) and DHFR$^{F31V}$-MTX (right) complexes. (A) Complex structures closest to the cluster center of a metastable bound state corresponding to the crystallographic pose (*i.e.* micro state index = 1 in Figs. 4 and 5). (B) Complex structures closest to the cluster center of a metastable bound state corresponding to the doorway state (*i.e.* micro state index = 5 in Fig. 4 and micro state index = 12 in Fig. 5). The protein is represented by a surface model, and F/V31 are colored in magenta. The ligand is depicted as orange sticks.

#### 2. Supplementary Tables

Table S1 Kinetic and thermodynamic parameters of inhibitor binding to DHFR with Asp27 in the deprotonated and protonated states.

The on-rate constant ($k_{on}$), off-rate constant ($k_{off}$), and binding free energy ($\Delta G$) were estimated for each model, as shown in Fig. 2. Values are presented as the mean ± SD of three independent MSM analyses. The off-rate constant for DHFR with Asp27 in the protonated state is described as $\log(k_{off})$ because of the larger computational errors arising from its calculation according to $k_{off} = k_{on} / \exp(-\Delta G/RT)$ (see Methods and Figure S6). The experimentally determined $k_{on}$, $k_{off}$, and $\Delta G$ values were retrieved from Table 1 of [1].

| **Model1** | Asp27 in the deprotonated state | | | $\Delta G$ change upon Asp27 protonation | Asp27 in the protonated state | | |
|---|---|---|---|---|---|---|---|
| | $k_{on}$ (1/Ms) | $\Delta G$ (kcal/mol) | $k_{off}$ (1/s) | | $k_{on}$ (1/Ms) | $\Delta G$ (kcal/mol) | $\log(k_{off}$ (1/s)) |
| DHFR$^{WT}$-MTX | $7.55 \pm 0.53 \times 10^6$ | -5.76 ± 0.04 | $2.85 \pm 0.05 \times 10^4$ | -6.27 ± 1.62 | $7.55 \pm 0.53 \times 10^6$ | -12.03 ± 1.62 | -1.83 ± 1.20 |
| DHFR$^{WT}$-DAM | $1.09 \pm 0.01 \times 10^7$ | -2.29 ± 0.04 | $7.21 \pm 0.31 \times 10^5$ | -3.44 ± 0.95 | $1.09 \pm 0.01 \times 10^7$ | -5.73 ± 0.95 | 2.89 ± 0.69 |
| DHFR$^{F31V}$-MTX | $2.84 \pm 0.31 \times 10^6$ | -3.78 ± 0.24 | $6.69 \pm 0.43 \times 10^4$ | -1.92 ± 2.79 | $2.84 \pm 0.31 \times 10^6$ | -5.70 ± 2.80 | 2.33 ± 2.07 |

| **Model2** | Asp27 in the deprotonated state | | | $\Delta G$ change upon Asp27 protonation | Asp27 in the protonated state | | |
|---|---|---|---|---|---|---|---|
| | $k_{on}$ (1/Ms) | $\Delta G$ (kcal/mol) | $k_{off}$ (1/s) | | $k_{on}$ (1/Ms) | $\Delta G$ (kcal/mol) | $\log(k_{off}$ (1/s)) |
| DHFR$^{WT}$-MTX | $3.68 \pm 0.59 \times 10^8$ | -6.82 ± 0.02 | $3.14 \pm 0.07 \times 10^4$ | -1.89 ± 1.69 | $3.68 \pm 0.59 \times 10^8$ | -8.71 ± 1.68 | 2.26 ± 1.29 |
| DHFR$^{WT}$-DAM | $2.53 \pm 0.05 \times 10^9$ | -4.08 ± 0.00 | $2.04 \pm 0.05 \times 10^6$ | 1.51 ± 0.35 | $2.53 \pm 0.05 \times 10^9$ | -2.57 ± 0.35 | 7.54 ± 0.26 |
| DHFR$^{F31V}$-MTX | $9.35 \pm 3.08 \times 10^8$ | -6.22 ± 0.01 | $8.82 \pm 0.42 \times 10^4$ | -0.23 ± 1.17 | $9.35 \pm 3.08 \times 10^8$ | -6.45 ± 1.17 | 4.30 ± 1.00 |

| **Model3** | Asp27 in the deprotonated state | | | $\Delta G$ change upon Asp27 protonation | Asp27 in the protonated state | | |
|---|---|---|---|---|---|---|---|
| | $k_{on}$ (1/Ms) | $\Delta G$ (kcal/mol) | $k_{off}$ (1/s) | | $k_{on}$ (1/Ms) | $\Delta G$ (kcal/mol) | $\log(k_{off}$ (1/s)) |
| DHFR$^{WT}$-MTX | $8.01 \pm 0.61 \times 10^6$ | -2.15 ± 0.02 | $1.30 \pm 0.02 \times 10^6$ | -4.79 ± 0.60 | $8.01 \pm 0.61 \times 10^6$ | -6.94 ± 0.60 | 1.88 ± 0.47 |
| DHFR$^{WT}$-DAM | $1.10 \pm 0.01 \times 10^7$ | -1.23 ± 0.04 | $9.92 \pm 0.60 \times 10^5$ | -5.21 ± 0.94 | $1.10 \pm 0.01 \times 10^7$ | -6.44 ± 0.93 | 2.38 ± 0.68 |
| DHFR$^{F31V}$-MTX | $2.89 \pm 0.32 \times 10^6$ | -0.68 ± 0.24 | $5.59 \pm 3.88 \times 10^5$ | -1.74 ± 2.59 | $2.89 \pm 0.32 \times 10^6$ | -2.42 ± 2.60 | 4.71 ± 1.93 |

| **Experimental** | $k_{on}$ (1/Ms) | $\Delta G$ (kcal/mol) | $k_{off}$ (1/s) |
|---|---|---|---|
| DHFR$^{WT}$-MTX | $3.6 \pm 0.2 \times 10^7$ | -8.9 ± 0.1 | $1.2 \pm 0.2 \times 10^1$ |
| DHFR$^{WT}$-DAM | $1.2 \pm 0.1 \times 10^7$ | -7.7 ± 0.1 | $3.5 \pm 0.5 \times 10^1$ |
| DHFR$^{F31V}$-MTX | $1.5 \pm 0.3 \times 10^5$ | -5.2 ± 0.4 | $2.6 \pm 1.0 \times 10^1$ |

Table S2 Free energy changes of the DHFR$^{WT}$-MTX complex upon protonation of Asp27. The values are presented as the mean ± SD of three independent FEP simulations. The snapshot closest to the cluster center of each metastable or unbound state was set as the initial structure of the free energy simulation. For metastable state index of 1, which corresponds to the crystallographic binding mode, multiple FEP simulations were performed using different initial structures to decrease the free-energy bias caused by insufficient sampling limited to around the initial structure.

| Trial of MSM analysis | Metastable state index | Free energy change (kcal/mol) |
|---|---|---|
| Trial 1 | 1 | 22.7 ± 0.6 |
| | 1 | 24.5 ± 0.3 |
| | 1 | 22.4 ± 0.8 |
| | 2 | 26.3 ± 0.3 |
| | 3 | 36.0 ± 0.4 |
| | Unbound state | 31.6 ± 2.1 |

| Trial of MSM analysis | Metastable state index | Free energy change (kcal/mol) |
|---|---|---|
| Trial 2 | 1 | 25.6 ± 0.9 |
| | 1 | 25.7 ± 0.2 |
| | 2 | 25.2 ± 1.4 |
| | 3 | 25.8 ± 0.4 |
| | 4 | 31.5 ± 0.5 |
| | 5 | 34.2 ± 1.0 |
| | Unbound state | 31.4 ± 1.3 |

| Trial of MSM analysis | Metastable state index | Free energy change (kcal/mol) |
|---|---|---|
| Trial 3 | 1 | 28.4 ± 1.1 |
| | 1 | 24.0 ± 1.0 |
| | 2 | 25.6 ± 2.0 |
| | 3 | 30.8 ± 0.3 |
| | 4 | 34.3 ± 0.8 |
| | 5 | 33.9 ± 0.8 |
| | 6 | 32.3 ± 0.7 |
| | 7 | 31.5 ± 1.4 |
| | 8 | 29.7 ± 0.5 |
| | 9 | 31.7 ± 0.4 |
| | 10 | 34.2 ± 0.2 |
| | 11 | 33.3 ± 0.5 |
| | Unbound state | 30.7 ± 1.6 |

Table S3 Free energy changes of the $DHFR^{WT}$-DAM complex upon protonation of Asp27. The values are presented as the mean ± SD of three independent free energy perturbation (FEP) simulations. The snapshot closest to the cluster center of each metastable or unbound state was set as the initial structure of the free energy simulation.

| Trial of MSM analysis | Metastable state index | Free energy change (kcal/mol) | Metastable state index | Free energy change (kcal/mol) |
|---|---|---|---|---|
| Trial 1 | 1 | 27.6 ± 0.2 | 21 | 34.1 ± 0.8 |
| | 2 | 33.6 ± 1.5 | 22 | 33.3 ± 1.1 |
| | 3 | 30.5 ± 1.7 | 23 | 32.3 ± 0.6 |
| | 4 | 32.5 ± 0.5 | Unbound state | 31.7 ± 1.5 |
| | 5 | 31.2 ± 0.6 | | |
| | 6 | 32.8 ± 0.6 | | |
| | 7 | 31.7 ± 0.8 | | |
| | 8 | 37.2 ± 1.6 | | |
| | 9 | 34.3 ± 0.3 | | |
| | 10 | 33.1 ± 1.4 | | |
| | 11 | 34.6 ± 1.5 | | |
| | 12 | 32.1 ± 0.3 | | |
| | 13 | 33.8 ± 0.4 | | |
| | 14 | 32.2 ± 1.1 | | |
| | 15 | 31.2 ± 1.5 | | |
| | 16 | 32.2 ± 0.5 | | |
| | 17 | 33.6 ± 2.1 | | |
| | 18 | 32.9 ± 0.5 | | |
| | 19 | 33.3 ± 0.9 | | |
| | 20 | 33.3 ± 0.3 | | |

| Trial of MSM analysis | Metastable state index | Free energy change (kcal/mol) |
|---|---|---|
| Trial 2 | 1 | 29.1 ± 1.0 |
| | 2 | 34.0 ± 1.4 |
| | 3 | 34.2 ± 0.2 |
| | 4 | 31.8 ± 0.5 |
| | 5 | 33.3 ± 0.4 |
| | 6 | 31.6 ± 1.0 |
| | 7 | 30.8 ± 1.1 |
| | 8 | 32.8 ± 0.7 |
| | 9 | 37.2 ± 0.9 |
| | 10 | 32.2 ± 0.4 |
| | 11 | 31.7 ± 0.9 |
| | 12 | 36.6 ± 1.7 |
| | 13 | 31.8 ± 0.6 |
| | 14 | 32.9 ± 1.0 |
| | 15 | 34.7 ± 0.1 |
| | 16 | 32.3 ± 1.8 |
| | 17 | 33.8 ± 0.8 |
| | 18 | 33.3 ± 1.1 |
| | Unbound state | 31.2 ± 0.3 |

| Trial of MSM analysis | Metastable state index | Free energy change (kcal/mol) |
|---|---|---|
| Trial 3 | 1 | 26.3 ± 0.7 |
| | 2 | 31.1 ± 2.1 |
| | 3 | 33.5 ± 0.4 |
| | 4 | 33.2 ± 1.0 |
| | 5 | 32.1 ± 1.4 |
| | 6 | 34.3 ± 0.8 |
| | 7 | 33.8 ± 0.6 |
| | Unbound state | 30.4 ± 0.5 |

Table S4 Free energy changes of the DHFR$^{F31V}$-MTX complex upon protonation of Asp27.

The values are presented as the mean ± SD of three independent FEP simulations. The snapshot closest to the cluster center of each metastable or unbound state was set as the initial structure of the free energy simulation. For metastable state index of 1, which corresponds to the crystallographic binding mode, multiple FEP simulations were performed using different initial structures to decrease the free-energy bias caused by insufficient sampling limited to around the initial structure.

| Trial of MSM analysis | Metastable state index | Free energy change (kcal/mol) | Metastable state index | Free energy change (kcal/mol) |
|---|---|---|---|---|
| Trial 1 | 1 | 23.7 ± 0.5 | 18 | 30.6 ± 0.8 |
| | 1 | 30.3 ± 1.6 | 19 | 30.1 ± 0.1 |
| | 1 | 32.5 ± 0.5 | 20 | 30.8 ± 0.9 |
| | 2 | 31.6 ± 0.2 | 21 | 30.8 ± 0.9 |
| | 3 | 40.1 ± 0.7 | 22 | 34.9 ± 1.7 |
| | 4 | 31.3 ± 0.6 | 23 | 29.4 ± 0.6 |
| | 5 | 29.7 ± 1.3 | 24 | 31.5 ± 0.2 |
| | 6 | 35.4 ± 1.5 | 25 | 32.8 ± 0.5 |
| | 7 | 27.6 ± 0.5 | 26 | 31.0 ± 1.7 |
| | 8 | 29.6 ± 1.6 | 27 | 32.7 ± 0.5 |
| | 9 | 32.0 ± 0.9 | 28 | 33.2 ± 0.8 |
| | 10 | 34.7 ± 0.6 | 29 | 31.0 ± 1.7 |
| | 11 | 31.9 ± 0.7 | 30 | 31.6 ± 0.5 |
| | 12 | 33.6 ± 0.7 | 31 | 30.3 ± 1.3 |
| | 13 | 31.2 ± 0.8 | 32 | 31.8 ± 0.4 |
| | 14 | 30.6 ± 0.8 | 33 | 31.5 ± 0.8 |
| | 15 | 36.0 ± 1.5 | 34 | 31.0 ± 0.4 |
| | 16 | 33.5 ± 0.9 | Unbound state | 30.6 ± 0.5 |
| | 17 | 31.7 ± 2.5 | | |

| Trial of MSM analysis | Metastable state index | Free energy change (kcal/mol) | Metastable state index | Free energy change (kcal/mol) |
|---|---|---|---|---|
| Trial 2 | 1 | 22.5 ± 0.5 | 17 | 28.5 ± 0.5 |
| | 1 | 33.3 ± 4.0 | 18 | 26.6 ± 1.5 |
| | 1 | 34.4 ± 0.3 | 19 | 40.3 ± 1.1 |
| | 1 | 20.2 ± 1.4 | 20 | 33.3 ± 2.3 |
| | 2 | 28.4 ± 0.9 | 21 | 33.4 ± 0.2 |
| | 3 | 30.7 ± 0.2 | 22 | 33.6 ± 0.2 |
| | 4 | 28.6 ± 3.4 | 23 | 32.2 ± 0.5 |
| | 5 | 28.9 ± 1.5 | 24 | 31.7 ± 0.8 |
| | 6 | 30.3 ± 0.7 | 25 | 33.0 ± 0.5 |
| | 7 | 30.7 ± 0.6 | Unbound state | 33.0 ± 0.8 |
| | 8 | 29.3 ± 0.2 | | |
| | 9 | 33.7 ± 0.9 | | |
| | 10 | 28.9 ± 0.1 | | |
| | 11 | 30.9 ± 0.6 | | |
| | 12 | 29.8 ± 0.6 | | |
| | 13 | 34.2 ± 1.5 | | |
| | 14 | 36.3 ± 2.5 | | |
| | 15 | 30.7 ± 0.7 | | |
| | 16 | 31.2 ± 0.9 | | |

| Trial of MSM analysis | Metastable state index | Free energy change (kcal/mol) | Metastable state index | Free energy change (kcal/mol) |
|---|---|---|---|---|
| Trial 3 | 1 | 32.0 ± 0.6 | 17 | 33.2 ± 0.6 |
| | 1 | 38.1 ± 1.1 | 18 | 30.8 ± 0.7 |
| | 1 | 40.1 ± 1.0 | 19 | 31.8 ± 0.4 |
| | 1 | 23.2 ± 0.3 | 20 | 31.6 ± 0.5 |
| | 2 | 26.9 ± 1.4 | Unbound state | 32.0 ± 2.0 |
| | 3 | 33.9 ± 0.9 | | |
| | 4 | 32.9 ± 0.4 | | |
| | 5 | 29.5 ± 0.9 | | |
| | 6 | 30.8 ± 0.4 | | |
| | 7 | 29.6 ± 0.5 | | |
| | 8 | 30.1 ± 0.4 | | |
| | 9 | 31.1 ± 1.2 | | |
| | 10 | 31.1 ± 0.3 | | |
| | 11 | 30.4 ± 0.3 | | |
| | 12 | 32.5 ± 0.4 | | |
| | 13 | 31.5 ± 1.4 | | |
| | 14 | 31.8 ± 0.6 | | |
| | 15 | 32.7 ± 1.4 | | |
| | 16 | 31.8 ± 0.3 | | |

Table S5 Contributions of each metastable state of the $DHFR^{WT}$-DAM complex in a network of fluxes from the unbound state to the active site-bound state. After all possible transition pathways linking microstates were computed by employing transition path theory (TPT), contribution of each metastable state was estimated by summing fluxes leaving (entering) all the microstates assigned to it. Metastable states that occupy more than 5% of the total flux are colored in red.

$DHFR^{WT}$-DAM

| Trial of MSM analysis | Metastable state index | Flux contribution | Metastable state index | Flux contribution |
|---|---|---|---|---|
| Trial 1 | 1 | $1.7\times10^{-3}$ | 19 | $2.2\times10^{-5}$ |
| | 2 | $1.8\times10^{-4}$ | 20 | $3.1\times10^{-4}$ |
| | 3 | $1.1\times10^{-4}$ | 21 | $1.8\times10^{-5}$ |
| | 4 | $2.0\times10^{-4}$ | 22 | $7.9\times10^{-5}$ |
| | 5 | $5.1\times10^{-4}$ | 23 | $2.5\times10^{-5}$ |
| | 6 | $1.2\times10^{-5}$ | | |
| | 7 | $3.0\times10^{-6}$ | | |
| | 8 | $8.0\times10^{-5}$ | | |
| | 9 | $1.3\times10^{-4}$ | | |
| | 10 | $1.6\times10^{-6}$ | | |
| | 11 | $9.6\times10^{-6}$ | | |
| | 12 | $2.8\times10^{-5}$ | | |
| | 13 | $3.3\times10^{-5}$ | | |
| | 14 | $4.4\times10^{-5}$ | | |
| | 15 | $2.6\times10^{-5}$ | | |
| | 16 | $6.5\times10^{-5}$ | | |
| | 17 | $9.5\times10^{-5}$ | | |
| | 18 | $2.0\times10^{-5}$ | | |

| Trial of MSM analysis | Metastable state index | Flux contribution |
|---|---|---|
| Trial 2 | 1 | $1.7\times10^{-3}$ |
| | 2 | $1.1\times10^{-4}$ |
| | 3 | $1.8\times10^{-4}$ |
| | 4 | $5.5\times10^{-5}$ |
| | 5 | $4.1\times10^{-5}$ |
| | 6 | $1.6\times10^{-4}$ |
| | 7 | $9.1\times10^{-5}$ |
| | 8 | $1.3\times10^{-4}$ |
| | 9 | $3.7\times10^{-5}$ |
| | 10 | $6.2\times10^{-5}$ |
| | 11 | $3.0\times10^{-4}$ |
| | 12 | $5.7\times10^{-6}$ |
| | 13 | $3.3\times10^{-5}$ |
| | 14 | $8.9\times10^{-6}$ |
| | 15 | $6.4\times10^{-4}$ |
| | 16 | $6.8\times10^{-5}$ |
| | 17 | $2.1\times10^{-5}$ |
| | 18 | $2.9\times10^{-5}$ |

| Trial of MSM analysis | Metastable state index | Flux contribution |
|---|---|---|
| Trial 3 | 1 | $1.7\times10^{-3}$ |
| | 2 | $8.2\times10^{-4}$ |
| | 3 | $1.1\times10^{-5}$ |
| | 4 | $4.2\times10^{-6}$ |
| | 5 | $1.4\times10^{-5}$ |
| | 6 | $8.2\times10^{-6}$ |
| | 7 | $7.8\times10^{-4}$ |

Table S6 Contributions of each metastable state of the $DHFR^{WT}$-MTX complex in a network of fluxes from the unbound state to the active site-bound state. After all possible transition pathways linking microstates were computed by employing TPT, contribution of each metastable state was estimated by summing fluxes leaving (entering) all the microstates assigned to it. Metastable states that occupy more than 5% of the total flux are colored in red.

$DHFR^{WT}$-MTX

| Trial of MSM analysis | Metastable state index | Flux contribution |
|---|---|---|
| Trial 1 | 1 | $5.2\times10^{-4}$ |
| | 2 | $2.7\times10^{-4}$ |
| | 3 | $5.8\times10^{-4}$ |

| Trial of MSM analysis | Metastable state index | Flux contribution |
|---|---|---|
| Trial 2 | 1 | $5.0\times10^{-4}$ |
| | 2 | $2.0\times10^{-4}$ |
| | 3 | $4.1\times10^{-5}$ |
| | 4 | $2.4\times10^{-5}$ |
| | 5 | $5.1\times10^{-4}$ |

| Trial of MSM analysis | Metastable state index | Flux contribution |
|---|---|---|
| Trial 3 | 1 | $5.1\times10^{-4}$ |
| | 2 | $2.1\times10^{-4}$ |
| | 3 | $4.2\times10^{-5}$ |
| | 4 | $1.3\times10^{-5}$ |
| | 5 | $6.9\times10^{-6}$ |
| | 6 | $2.7\times10^{-6}$ |
| | 7 | $1.2\times10^{-4}$ |
| | 8 | $2.9\times10^{-5}$ |
| | 9 | $8.3\times10^{-6}$ |
| | 10 | $4.3\times10^{-4}$ |
| | 11 | $1.2\times10^{-5}$ |

Table S7 Contributions of each metastable state of the DHFR$^{F31V}$-MTX complex in a network of fluxes from the unbound state to the active site-bound state. After all possible transition pathways linking microstates were computed by employing TPT, contribution of each metastable state was estimated by summing fluxes leaving (entering) all the microstates assigned to it. Metastable states that occupy more than 5% of the total flux are colored in red.

DHFR$^{F31V}$-MTX

| Trial of MSM analysis | Metastable state index | Flux contribution | Metastable state index | Flux contribution |
|---|---|---|---|---|
| Trial 1 | 1 | $3.8 \times 10^{-4}$ | 21 | $3.4 \times 10^{-5}$ |
| | 2 | $7.4 \times 10^{-5}$ | 22 | $2.1 \times 10^{-6}$ |
| | 3 | $8.8 \times 10^{-6}$ | 23 | $3.5 \times 10^{-5}$ |
| | 4 | $2.8 \times 10^{-5}$ | 24 | $7.7 \times 10^{-6}$ |
| | 5 | $1.5 \times 10^{-5}$ | 25 | $2.1 \times 10^{-5}$ |
| | 6 | $4{,}7 \times 10^{-6}$ | 26 | $2.5 \times 10^{-5}$ |
| | 7 | $1.5 \times 10^{-5}$ | 27 | $1.3 \times 10^{-7}$ |
| | 8 | $6.7 \times 10^{-6}$ | 28 | $3.0 \times 10^{-5}$ |
| | 9 | $5.2 \times 10^{-5}$ | 29 | $4.7 \times 10^{-6}$ |
| | 10 | $3.4 \times 10^{-5}$ | 30 | $2.9 \times 10^{-6}$ |
| | 11 | $1.1 \times 10^{-5}$ | 31 | $3.8 \times 10^{-6}$ |
| | 12 | $1.2 \times 10^{-5}$ | 32 | $6.3 \times 10^{-7}$ |
| | 13 | $1.5 \times 10^{-5}$ | 33 | $8.9 \times 10^{-7}$ |
| | 14 | $7.1 \times 10^{-5}$ | 34 | $2.2 \times 10^{-6}$ |
| | 15 | $2.5 \times 10^{-5}$ | | |
| | 16 | $1.8 \times 10^{-5}$ | | |
| | 17 | $7.1 \times 10^{-5}$ | | |
| | 18 | $3.4 \times 10^{-5}$ | | |
| | 19 | $1.4 \times 10^{-6}$ | | |
| | 20 | $2.2 \times 10^{-4}$ | | |

| Trial of MSM analysis | Metastable state index | Flux contribution | Metastable state index | Flux contribution |
|---|---|---|---|---|
| Trial 2 | 1 | $3.5 \times 10^{-4}$ | 21 | $1.8 \times 10^{-5}$ |
| | 2 | $2.6 \times 10^{-5}$ | 22 | $1.6 \times 10^{-5}$ |
| | 3 | $2.7 \times 10^{-5}$ | 23 | $8.0 \times 10^{-6}$ |
| | 4 | $9.6 \times 10^{-6}$ | 24 | $3.9 \times 10^{-6}$ |
| | 5 | $2.6 \times 10^{-6}$ | 25 | $1.5 \times 10^{-6}$ |
| | 6 | $7{,}7 \times 10^{-6}$ | | |
| | 7 | $6.9 \times 10^{-6}$ | | |
| | 8 | $5.1 \times 10^{-6}$ | | |
| | 9 | $2.0 \times 10^{-5}$ | | |
| | 10 | $9.5 \times 10^{-6}$ | | |
| | 11 | $1.7 \times 10^{-5}$ | | |
| | 12 | $1.0 \times 10^{-4}$ | | |
| | 13 | $4.3 \times 10^{-6}$ | | |
| | 14 | $1.9 \times 10^{-5}$ | | |
| | 15 | $1.5 \times 10^{-5}$ | | |
| | 16 | $1.2 \times 10^{-4}$ | | |
| | 17 | $4.4 \times 10^{-5}$ | | |
| | 18 | $1.4 \times 10^{-4}$ | | |
| | 19 | $3.8 \times 10^{-6}$ | | |
| | 20 | $2.3 \times 10^{-5}$ | | |

| Trial of MSM analysis | Metastable state index | Flux contribution |
|---|---|---|
| Trial 3 | 1 | $4.4 \times 10^{-4}$ |
| | 2 | $2.2 \times 10^{-6}$ |
| | 3 | $4.2 \times 10^{-6}$ |
| | 4 | $1.2 \times 10^{-5}$ |
| | 5 | $3.3 \times 10^{-6}$ |
| | 6 | $2.1 \times 10^{-4}$ |
| | 7 | $2.9 \times 10^{-4}$ |
| | 8 | $1.6 \times 10^{-5}$ |
| | 9 | $1.0 \times 10^{-4}$ |
| | 10 | $1.0 \times 10^{-5}$ |
| | 11 | $3.5 \times 10^{-5}$ |
| | 12 | $4.7 \times 10^{-5}$ |
| | 13 | $1.3 \times 10^{-5}$ |
| | 14 | $7.1 \times 10^{-7}$ |
| | 15 | $1.6 \times 10^{-5}$ |
| | 16 | $6.1 \times 10^{-6}$ |
| | 17 | $5.2 \times 10^{-6}$ |
| | 18 | $6.0 \times 10^{-7}$ |
| | 19 | $3.3 \times 10^{-6}$ |
| | 20 | $3.1 \times 10^{-6}$ |

Table S8 $DHFR^{WT}$-MTX, $DHFR^{WT}$-DAM, and $DHFR^{F31V}$-MTX binding free energies ($\Delta G$) estimated by the MP-CAFEE method [3]. Electrostatic (coulomb) and van der Waals (vdW) contributions in $\Delta G$ values are also indicated. Representative five micro states assigned to a metastable state corresponding to the crystallographic binding mode (*i.e.* metastable state index = 1 in Fig. 3-5) were extracted. A snapshot closest to the cluster center of each micro state was set to the initial structure of the free energy simulation. $\Delta G$ was computed according to a protocol described in the previous study [4]. Data are presented as the mean ± SD of five independent free energy simulations started from these initial structures.

| | ΔG | coulomb | vdW |
|---|---|---|---|
| **$DHFR^{WT}$-MTX** | **-27.71 ± 5.26** | **-18.75 ± 5.37** | **-8.96 ± 0.65** |
| **$DHFR^{WT}$-DAM** | **-6.52 ± 1.79** | **-2.52 ± 1.50** | **-4.00 ± 0.80** |
| **$DHFR^{F31V}$-MTX** | **-19.14 ± 3.23** | **-8.74 ± 3.87** | **-10.4 ± 1.29** |

**kcal/mol**

#### 3. Supplementary References


1. Taira, K.; Benkovic, S. J., Evaluation of the importance of hydrophobic interactions in drug binding to dihydrofolate reductase. *Journal of medicinal chemistry* **1988,** *31* (1), 129-37.
2. Ono, F.; Chiba, S.; Isaka, Y.; Matsumoto, S.; Ma, B.; Katayama, R.; Araki, M.; Okuno, Y., Improvement in predicting drug sensitivity changes associated with protein mutations using a molecular dynamics based alchemical mutation method. *Scientific reports* **2020,** *10* (1), 2161.
3. Fujitani, H.; Tanida, Y.; Matsuura, A., Massively parallel computation of absolute binding free energy with well-equilibrated states. *Phys. Rev. E* **2009,** *79* (2), 021914.
4. Araki, M.; Kamiya, N.; Sato, M.; Nakatsui, M.; Hirokawa, T.; Okuno, Y., The Effect of Conformational Flexibility on Binding Free Energy Estimation between Kinases and Their Inhibitors. *Journal of chemical information and modeling* **2016,** *56* (12), 2445-2456.
